# DeepUMQA3: a web server for model quality assessment of protein complexes

**DOI:** 10.1101/2023.04.24.538194

**Authors:** Jun Liu, Dong Liu, Guijun Zhang

## Abstract

Model quality assessment is a crucial part of protein structure prediction and a gateway to proper usage of models in biomedical applications. Many methods have been proposed for assessing the quality of structural models of protein monomers, but few methods for evaluating protein complex models. As protein complex structure prediction becomes a new challenge, model quality assessment methods that can provide accurate evaluation of complex structures are urgently required. Here, we present DeepUMQA3, a web server for evaluating protein complex structures using deep neural network. For an input complex structure, features are extracted from three levels of overall complex, intra-monomer, and inter-monomer, and a improved deep residual neural network is used to predict per-residue lDDT and interface residue accuracy. DeepUMQA3 ranks first in the blind test of interface residue accuracy estimation in CASP15, with Pearson, Spearman and AUC of 0.564, 0.535 and 0.755 under the lDDT measurement, which are 18.5%, 23.6% and 10.9% higher than the second-best method, respectively. DeepUMQA3 can also accurately assess the accuracy of all residues in the entire complex and distinguish high- and low-precision residues/models. The websever of DeepUMQA3 are freely available at http://zhanglab-bioinf.com/DeepUMQA_server/.

## 1 Introduction

Predicting the three-dimensional structure of proteins is one of the major basic research issues in the field of bioinformatics, and it is important for understanding protein functions, innovative drug development, and disease treatment (Senior et al., 2020; Baek et al., 2021). Structural prediction methods typically generate many alternative models, which are then evaluated using model quality assessment procedures to select the best model and/or further guide model refinement (Liu et al., 2020; Zhou et al., 2022). Estimation of model accuracy has been an independent prediction category for CASP since 2007 (Kwon et al., 2021). According to the number of models used, model quality assessment (MQA) or estimation of model accuracy (EMA) methods can be divided into single-model methods (Uziela et al., 2017; Cao et al., 2017; Olechnovič and Venclovas, 2017; Shuvo et al., 2020) and multi-model (consensus) methods (Cheng et al., 2019; McGuffin et al., 2021; Ye et al., 2021). The multi-model methods usually take a model pool as the input and use information from other protein models in the model pool to evaluate the accuracy of the current model, and its performance depends largely on the number and diversity of protein models in the input model pool. The single model methods directly evaluate a single model without dependence on other models, so it has received more and more attention and research (Pagès et al., 2019; Baldassarre et al., 2021; Hiranuma et al., 2021).

We previously proposed single-model method, DeepUMQA (Guo et al., 2022), designed a residue-level USR to characterize the relationship between residues and protein topological structures, and using convolutional neural networks to predict the accuracy of each residue. DeepUMQA2 further integrates co-evolution-based sequence features and template-based structural features to complement the shared properties between different models of the same protein, and uses an improved neural network to predict the accuracy of local residues (Liu et al., 2023). DeepUMQA2 outperforms DeepUMQA by more than 15% and is able to select more accurate models for state-of-the-art structure prediction methods such as AlphaFold2 (Jumper et al., 2021). The DeepUMQA and DeepUMQA2 have been among the best quality assessors in CAMEO since their introduction. Although the previous version of DeepUMQA has made progress on model quality assessment of protein monomers, they were unable to assess the structure of protein complexes. As the protein monomer structure prediction problem has been largely solved, the structure prediction of protein complexes has become the new challenge. Therefore, methods that can accurately assess the structural quality of complexes are needed (Kwon et al., 2021). In CASP15, the EMA category was changed from protein monomer accuracy estimation to protein complex accuracy estimation.

In this work, we proposed DeepUMQA3, a web server for protein complex model quality assessment. On the basis of DeepUMQA and DeepUMQA2, new features were designed for complex structures, and the lDDT of each residue and the accuracy of interface residues were predicted using an improved deep neural network. DeepUMQA3 ranked first in the accuracy estimation of protein complex interface residues in CASP15, and the web server of DeepUMQA3 provides fast and accurate interface residue accuracy prediction and per-residue lDDT prediction services for protein complexes.

## 2 Methods

The flowchart of DeepUMQA3 is shown in Figure 1. For the complex structure to be evaluated, DeepUMQA3 describes it from three levels: overall complex features, intra-monomer features, and inter-monomer features. At the level of overall complex, the overall complex is regarded as a large monomer structure. Considering that protein complexes are discontinuous in sequence, features independent of residue order were extracted, including overall USR, residue voxelization (Pagès et al., 2019), inter-residue distance and orientations (Yang et al., 2020), and amino acid properties (Henikoff and Henikoff, 1992; Meiler et al., 2001). At the level of intra-monomer, the features of each monomer are extracted separately, including the sequence embedding generated by ESM-1b (Lin et al., 2023), secondary structure (Kabsch and Sander, 1983), and Rosetta energy terms (Leaver-Fay et al., 2011). At the inter-monomer level, the attention map of the inter-monomer paired sequence (Lin et al., 2023) was used to describe the sequence relationship between monomers. In addition, inter-monomer USR is designed to describe the relationship between residues in one monomer and topologies of other monomers. The features of these three levels were fed into a deep convolutional neural network coupled with triangular update and axial attention to predict the inter-residue distance deviation and the inter-residue contact map with a threshold of 15Å to calculate per-residue lDDT and interface residue accuracy. A detailed description of the method can be found in the invited paper for CASP15 (Liu et al., 2023).

**Fig. 1.**
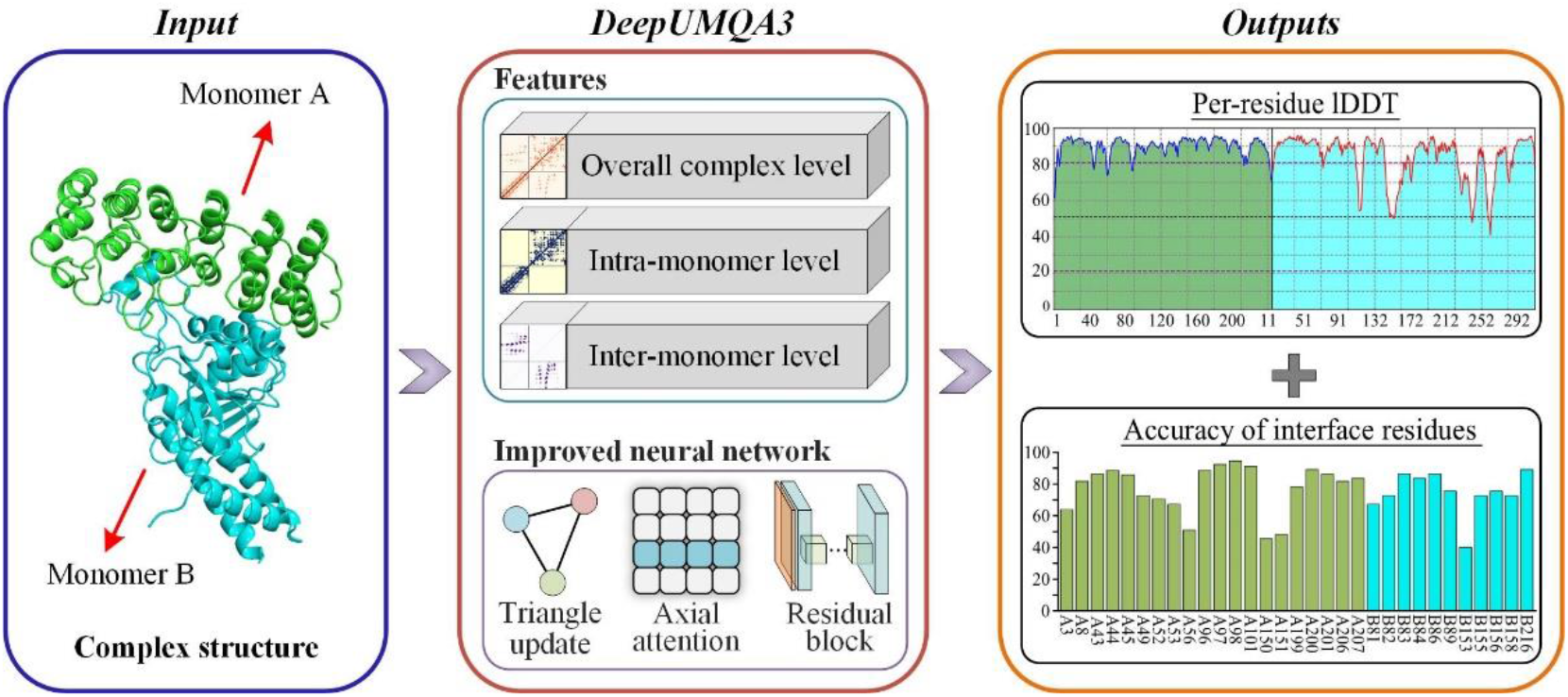
The flowchart of DeepUMQA3. For the input complex structure, it is described from three aspects: overall complex features, intra-monomer features, and inter-monomer features. Then, the extracted features are fed into a residual neural network coupled with triangle update and axial attention to predict the lDDT of each residue and the accuracy of the interface residues.

## 3 Results

### 3.1 Performance of DeepUMQA3

DeepUMQA3 (Group name: GuijunLab-RocketX) participated in the blind test of the EMA of CASP15 and ranked first in the accuracy estimation of interface residues (see Supplementary Figure S1). The performance of the participating methods of interface residue accuracy estimation in CASP15 is shown in Table 1. DeepUMQA3’s Pearson correlation coefficient (PCC), Spearman correlation coefficient (SCC), and the area under the receiver operating characteristic curve (AUC) on lDDT (Mariani et al., 2013) and CAD (Olechnovič et al., 2013) measurements are much better than those of other participating methods (see Supplementary Figure S2). For the measurement of lDDT, DeepUMQA3’s PCC (0.564) and SCC (0.535) were 18.5% and 23.6% higher than the second ranked methods, respectively, which was the only method exceeding 0.5. The AUC (0.755) of DeepUMQA3 is 10.9% higher than that of the second ranked method, indicating that DeepUMQA3 is easier to distinguish between high- and low-precision interface residues. DeepUMQA3 achieved the best performance (with highest PCC of lDDT) on 21 out of 36 submitted targets (3 were missed when submitted). Evaluation results on the missing 3 targets using the programs provided by the assessor show that DeepUMQA3 outperforms other methods on 2 of them (see Supplementary Figure S3). The performance of DeepUMQA3 on both of homomers and heteromers are also the best among all methods (see Supplementary Table S1). DeepUMQA3 achieved the best performance on 4 out of 5 nanobody complex targets and all 3 antibody-antigen complex targets, the best among all methods (see Supplementary Table S2 and Figure S3).

**Table 1.** The performance of different model quality assessment methods in the interface residue accuracy estimation blind test in CASP15.

| Method | IDDT |  |  | CAD |  |  |
| --- | --- | --- | --- | --- | --- | --- |
|  | Pearson | Spearman | AUC | Pearson | Spearman | AUC |
| **DeepUMQA3** | **0.564** | **0.535** | **0.755** | **0.505** | **0.456** | **0.714** |
| ModFOLDdockS | 0.455 | 0.416 | 0.674 | 0.42 | 0.379 | 0.66 |
| ModFOLDdockR | 0.476 | 0.433 | 0.681 | 0.411 | 0.369 | 0.651 |
| VoroIF | 0.333 | 0.339 | 0.664 | 0.272 | 0.271 | 0.619 |
| Venclovas | 0.332 | 0.338 | 0.664 | 0.271 | 0.271 | 0.619 |
| FoldEver | 0.277 | 0.279 | 0.625 | 0.217 | 0.194 | 0.583 |
| ModFOLDdock | 0.243 | 0.227 | 0.584 | 0.209 | 0.2 | 0.572 |
| APOLLO | 0.192 | 0.213 | 0.565 | 0.156 | 0.159 | 0.549 |
| MULTICOM_deep | 0.091 | 0.094 | 0.538 | 0.082 | 0.091 | 0.534 |
| Manifold | 0.18 | 0.176 | 0.542 | 0.153 | 0.149 | 0.529 |
| DLA-Ranker | 0.1 | 0.112 | 0.529 | 0.093 | 0.095 | 0.526 |
| MASS | 0.151 | 0.172 | 0.527 | 0.141 | 0.152 | 0.521 |
| LAW | 0.169 | 0.168 | 0.525 | 0.143 | 0.133 | 0.513 |

Note: Pearson and Spearman indicated the correlation between the accuracy of predicted interface residues and the real IDDT/CAD of interface residues. AUC is used to measure the ability of the model quality assessment method to distinguish high-and low-precision interface residues.

DeepUMQA3 can not only accurately assess the accuracy of interface residues, but also accurately predict the accuracy of all residues in the entire complex, and can distinguish high-precision and low-precision residues/models, which will provide important information for future structure refinement (see Supplementary Table S3, S4 and Figure S4). Supplementary Figure S5 presents an example of DeepUMQA3 evaluating model accuracy on one structural model of the target T1170. It can be found that the predicted lDDT of the interface residues is very close to the real one, and the PCC, average residue-wise S-score error (ASE) (Kwon et al., 2021) and AUC are 0.771, 0.923 and 0.872, respectively. For all residues in the entire complex, the predicted lDDT can accurately capture the changing of residue accuracy, and can easily distinguish high-quality and low-quality regions from the overall structure.

### 3.2 The web server of DeepUMQA3

The web server of DeepUMQA3 is freely available at http://zhanglab-bioinf.com/DeepUMQA_server/. The only mandatory input is the query protein complex structure information in PDB format (see Supplementary Figure S6). The web server allows users to input up to 10 complex structures at a time for evaluation. For the complex structures submitted by users, DeepUMQA3 first extracts features, then utilizes pre-trained network models to predict the accuracy of per-residue lDDT and interface residues, and finally generates a result page. If the user provides an email address, they will receive a task confirmation email after submitting the task and an email with evaluation results and web page when the task is completed. Supplementary Figure S7 demonstrates an example result web page. For each complex structure submitted by the user, the result web page will display the 3D structure, a graph of the per-residue lDDT, and the accuracy of the interface residues for each chain. Users can download each result individually, or download a compressed package of all results.

## Supplementary Information

### Supplementary Tables

**Table S1.** Performance of all methods under lDDT measurement for 19 homomer targets and 20 heteromer targets in the accuracy estimation of interface residues in CASP15.

| Type | Homomers (19) |  |  | Heteromers (20) |  |  |
| --- | --- | --- | --- | --- | --- | --- |
|  | Pearson | Spearman | AUC | Pearson | Spearman | AUC |
| DeepUMQA3 | 0.591 | 0.538 | 0.758 | 0.550 | 0.536 | 0.753 |
| ModFOLDdockR | 0.517 | 0.466 | 0.698 | 0.436 | 0.401 | 0.664 |
| ModFOLDdockS | 0.448 | 0.404 | 0.669 | 0.461 | 0.428 | 0.678 |
| ModFOLDdock | 0.276 | 0.256 | 0.596 | 0.211 | 0.200 | 0.572 |
| Manifold | 0.181 | 0.155 | 0.531 | 0.179 | 0.195 | 0.551 |
| MULTICOM_deep | 0.086 | 0.090 | 0.541 | 0.096 | 0.098 | 0.535 |
| Venclovas | 0.341 | 0.336 | 0.665 | 0.322 | 0.341 | 0.662 |
| MASS | 0.130 | 0.162 | 0.529 | 0.175 | 0.182 | 0.525 |
| LAW | 0.114 | 0.123 | 0.539 | 0.235 | 0.221 | 0.509 |
| APOLLO | 0.228 | 0.255 | 0.587 | 0.150 | 0.163 | 0.538 |
| VoroIF | 0.343 | 0.337 | 0.665 | 0.322 | 0.341 | 0.662 |
| FoldEver | 0.322 | 0.322 | 0.642 | 0.229 | 0.234 | 0.607 |
| DLA-Ranker | 0.095 | 0.099 | 0.533 | 0.105 | 0.124 | 0.525 |

**Table S2.** Performance of all methods under lDDT measurement for 5 nanobody complex targets and 3 antibody-antigen complex targets in the accuracy estimation of interface residues in CASP15.

| Type | Nanobody targets (5) |  |  | Antibody-antigen targets (3) |  |  |
| --- | --- | --- | --- | --- | --- | --- |
|  | Pearson | Spearman | AUC | Pearson | Spearman | AUC |
| DeepUMQA3 | 0.493 | 0.489 | 0.723 | 0.592 | 0.667 | 0.824 |
| ModFOLDdockR | 0.377 | 0.33 | 0.619 | 0.338 | 0.402 | 0.699 |
| ModFOLDdockS | 0.267 | 0.231 | 0.581 | 0.374 | 0.51 | 0.736 |
| ModFOLDdock | 0.215 | 0.229 | 0.586 | 0.182 | 0.235 | 0.623 |
| Manifold | 0.197 | 0.22 | 0.528 | 0.128 | 0.186 | 0.571 |
| MULTICOM_deep | 0.172 | 0.197 | 0.616 | 0.038 | 0.012 | 0.5 |
| Venclovas | 0.103 | 0.116 | 0.549 | NA | NA | NA |
| MASS | 0.12 | 0.136 | 0.562 | 0.445 | 0.417 | 0.5 |
| LAW | 0.438 | 0.435 | 0.5 | 0.158 | 0.084 | 0.5 |
| APOLLO | 0.137 | 0.175 | 0.5 | 0.289 | 0.356 | 0.663 |
| VoroIF | 0.103 | 0.116 | 0.549 | NA | NA | NA |
| FoldEver | 0.206 | 0.21 | 0.595 | 0.245 | 0.274 | 0.637 |
| DLA-Ranker | 0.08 | 0.085 | 0.524 | 0.215 | 0.253 | 0.5 |

**Table S3.** Performance of DeepUMQA3 for predicting per-residue lDDT of the overall complex on different type of targets in CASP15 (Local QA level).

| Type | Pearson | Spearman | Kendall | AUC | MAE |
| --- | --- | --- | --- | --- | --- |
| All (39) | 0.625 | 0.551 | 0.395 | 0.823 | 0.138 |
| Homomer (19) | 0.617 | 0.555 | 0.401 | 0.818 | 0.157 |
| Heteromer (20) | 0.631 | 0.547 | 0.389 | 0.828 | 0.120 |
| Nanobody (5) | 0.555 | 0.476 | 0.331 | 0.789 | 0.122 |
| Antibody-antigen (3) | 0.700 | 0.598 | 0.426 | 0.875 | 0.087 |

**Note:** Local QA is calculated based on the predicted and real IDDT for each residue. Pearson, Spearman, and Kendall are used to measure the correlation of the predicted residue's IDDT with the real IDDT. AUC is used to measure the ability of the predicted IDDT to discriminate high-/low-precision residues. MAE was used to measure the difference between the predicted residue IDDT and the real IDDT.

**Table S4.**
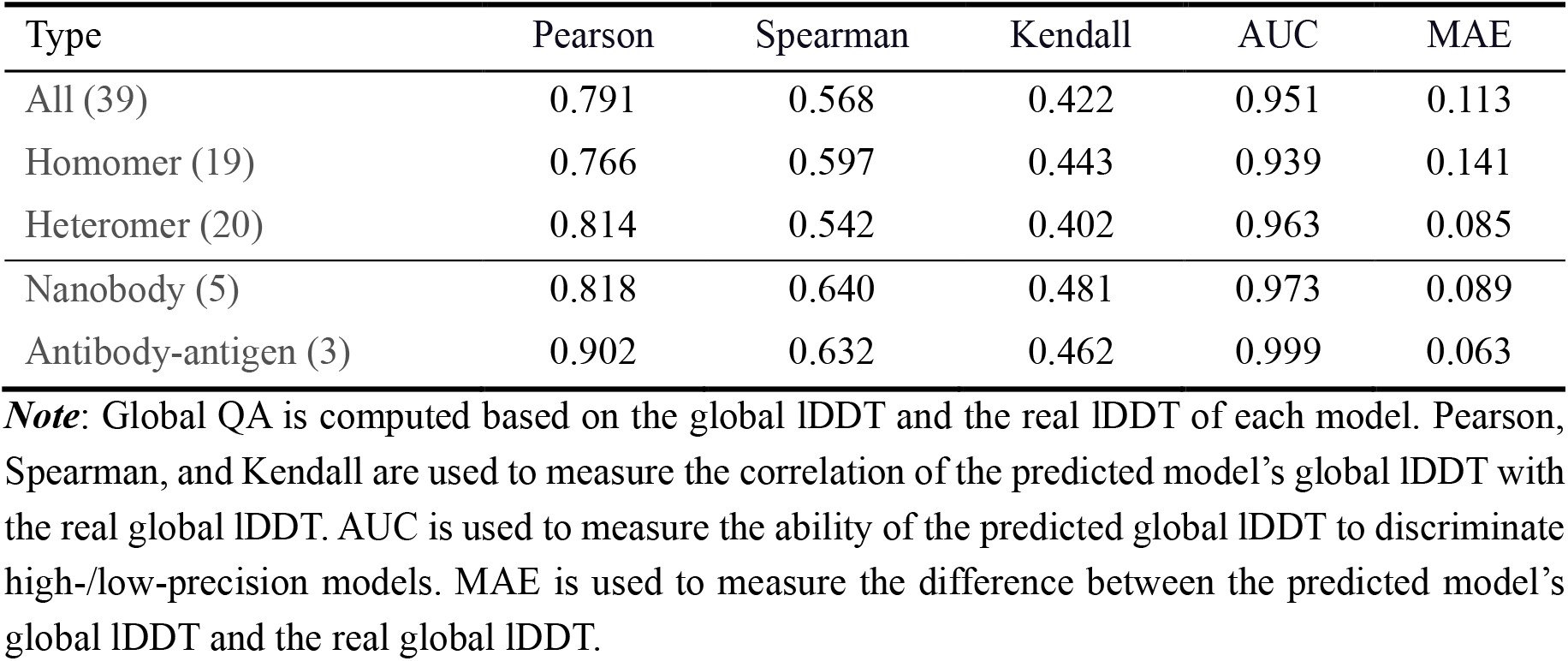
Performance of DeepUMQA3 for predicting per-residue lDDT of the overall complex on different type of targets in CASP15 (Global QA level).

### Supplementary Figures

**Figure S1.**
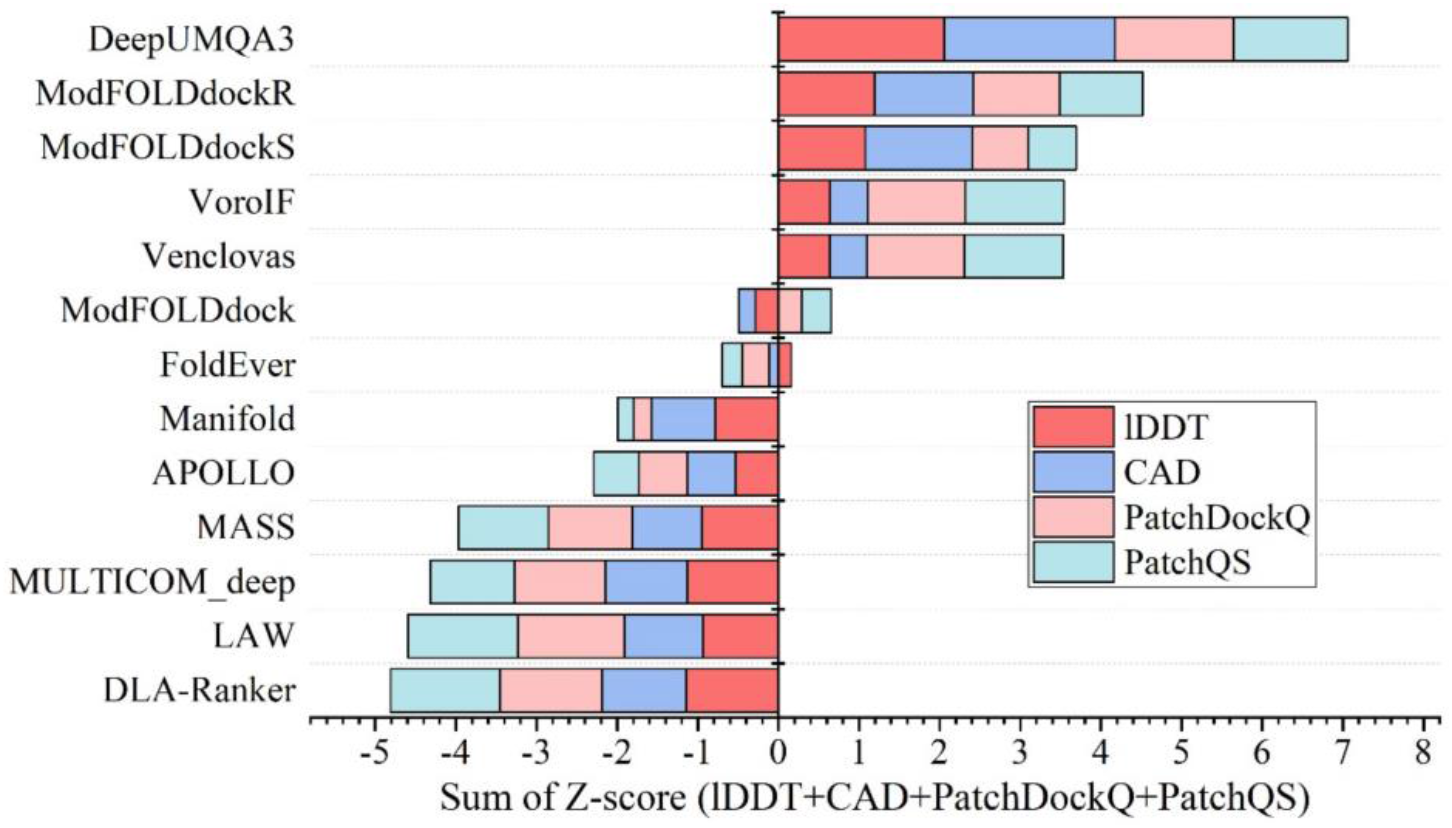
Ranking of the methods for interface residue precision estimation in CASP15 according to the sum of average Z-scores for lDDT (Red), CAD (blue), PatchDockQ (light red) and PatchQS(cyan). The Z-score of each item is weighted according to the Z-score of Pearson, Spearman and AUC according to the weight of 0.1:0.5:1. The data comes from the CASP15 official website (https://predictioncenter.org/casp15/qa_local.cgi). The group name of DeepUMQA3 in CASP15 is “GuijunLab-RocketX”.

**Figure S2.**
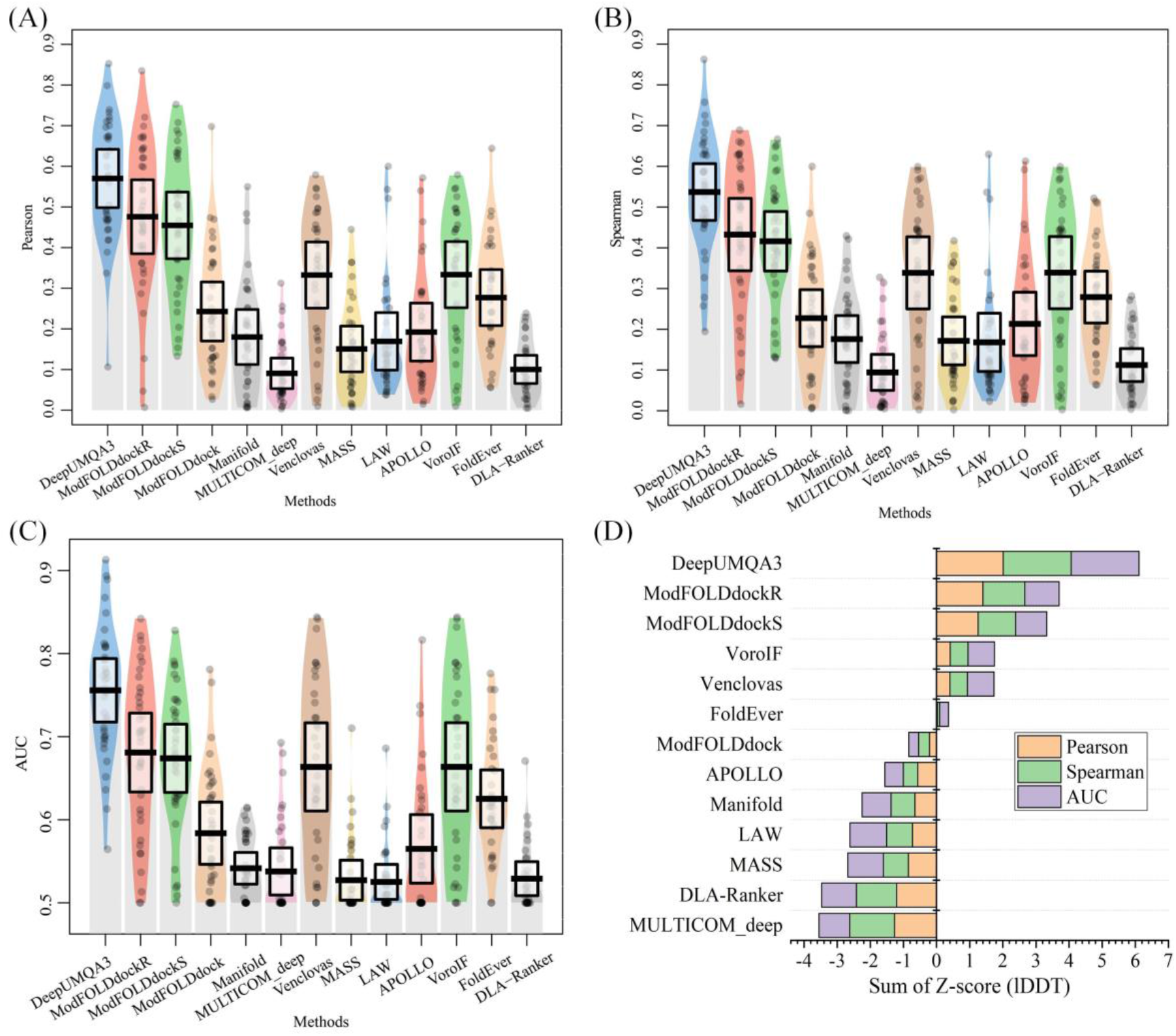
Performance of methods for assessing the accuracy of interface residues under lDDT measurement in CASP15. (A), (B), and (C) are the pirate graphs of Pearson, Spearman, and AUC for all participating methods on 39 targets, respectively. (D) is the ranking of Z-socre of Pearson, Spearman and AUC.

**Figure S3.**
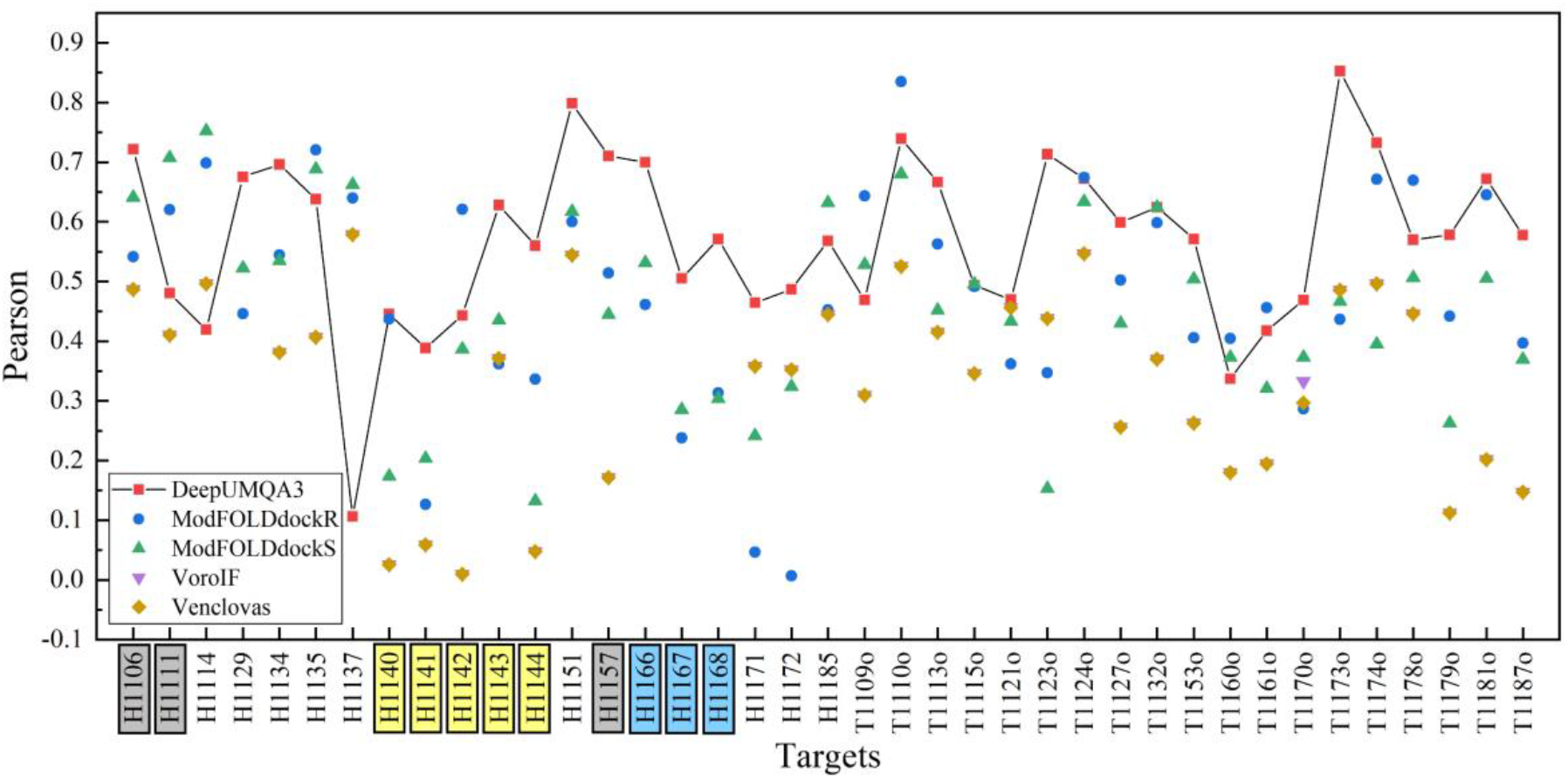
The Pearson correlation coefficient under lDDT measurement of the top 5 methods for interface residue accuracy evaluation on the 39 targets in CASP15. The targets in the gray boxes are the targets that were missed when DeepUMQA3 was submitted the results, and we evaluated them using the scripts provided by the reviewers. The target in the yellow box is the nanobody complex, and the target in the blue box is the antibody-antigen complex.

**Figure S4.**
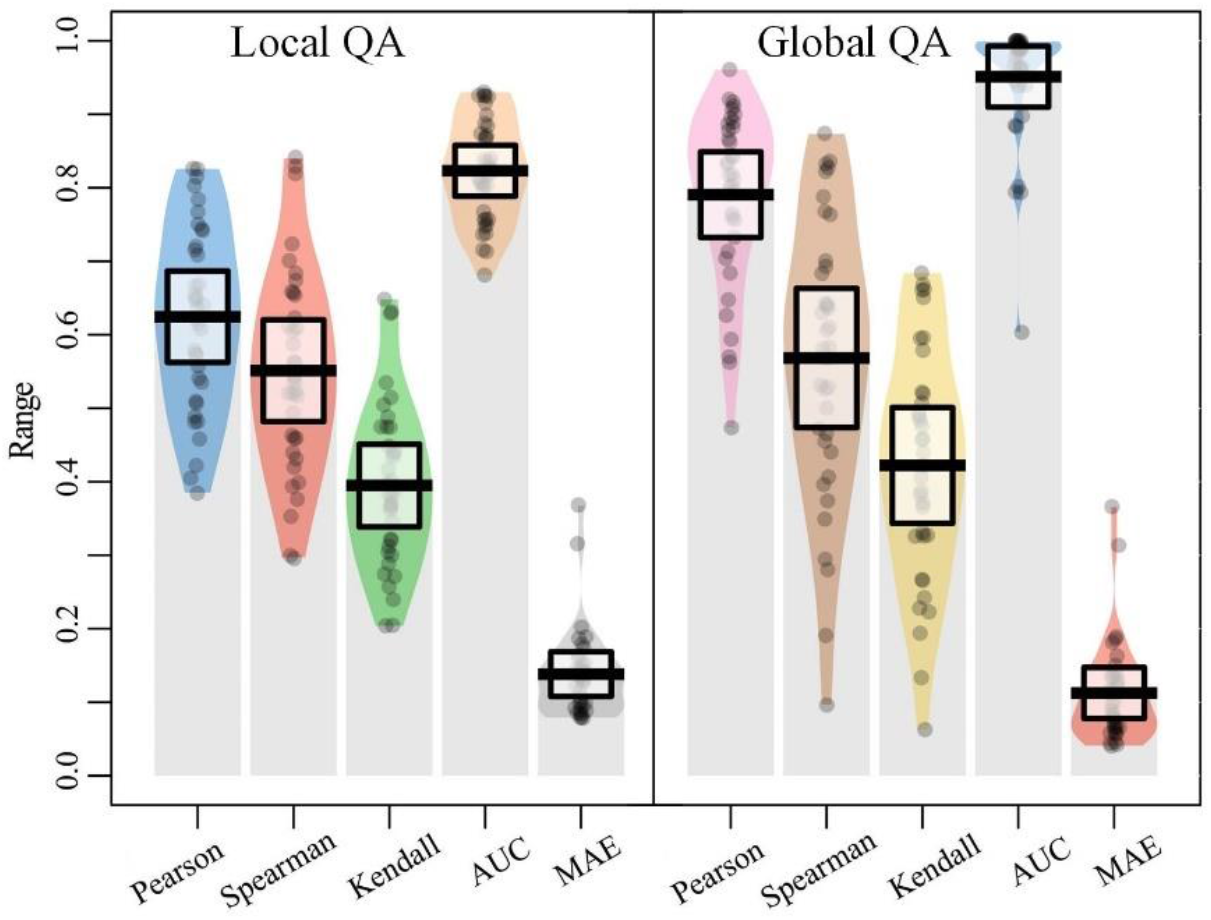
Pirate graphs on different performance indicators of DeepUMQA3 for predicting per-residue lDDT of overall complex in CASP15. On the left is the pirate map of Local QA, and on the right is the pirate map of Global QA. The higher the Pearson, Spearman, and Kendall, the stronger the correlation between the predicted lDDT and the real lDDT. The higher the AUC, the stronger the ability of DeepUMQA3 to distinguish high-/low-precision residues/models. The smaller the MAE, the difference in lDDT is smaller.

**Figure S5.**
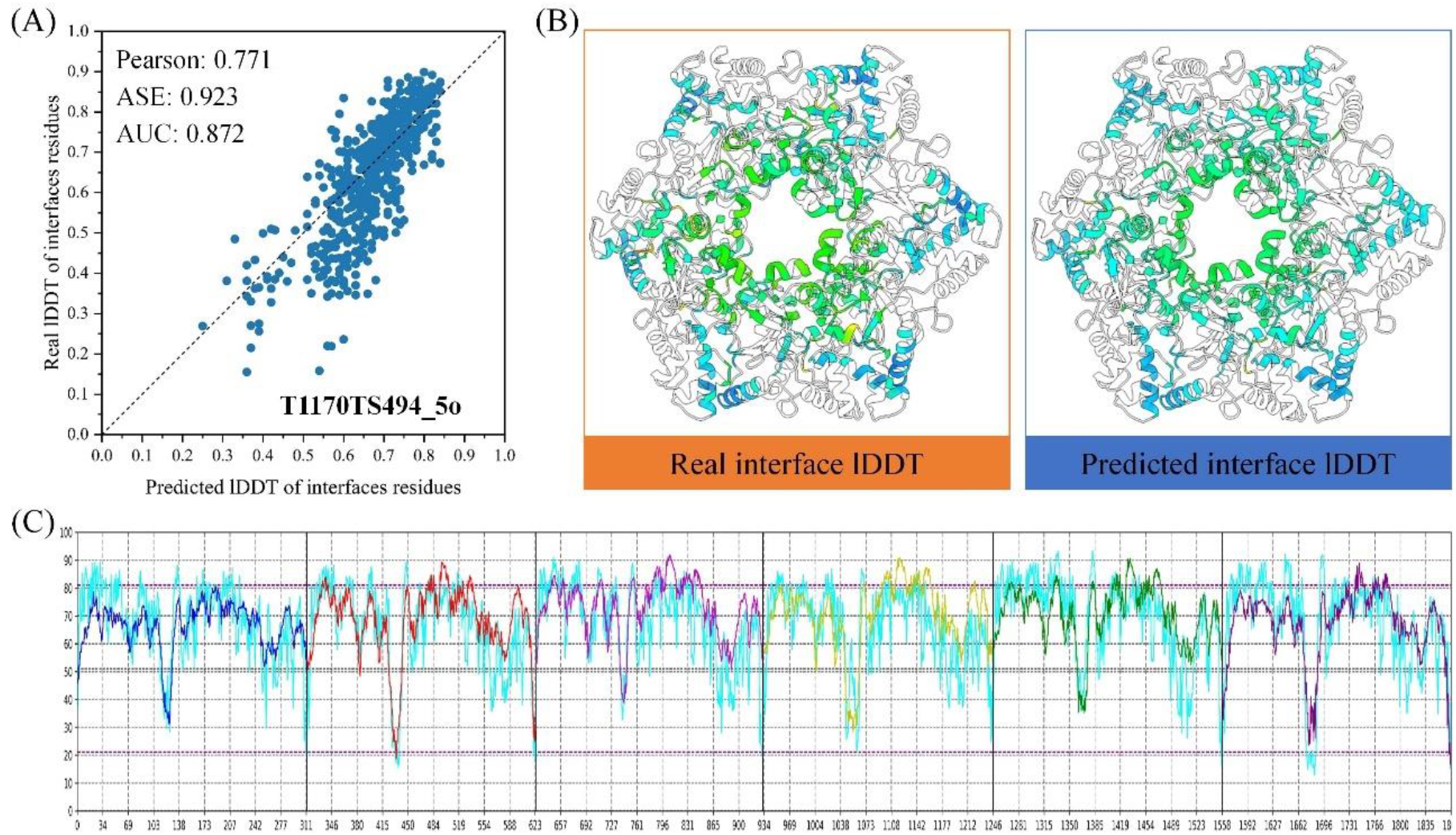
An example of DeepUMQA3 evaluating the model on the structural model T1170TS494_5o of the target T1170. T1170 is a homomer composed of 6 identical monomer structures, containing 1908 residues. (A) Head-to-head comparison of the predicted interface residue lDDT with the real lDDT. (B) The real interface residue lDDT (left) and the predicted interface residue lDDT (right) in the structural model. The colored parts represent the interface residues, and red to blue represents lDDT from 0 to 100. (C) Real lDDT (cyan) and predicted lDDT (other colors) for all residues in the overall protein complex, with predicted lDDT for different monomers indicated in different colors.

**Figure S6.**
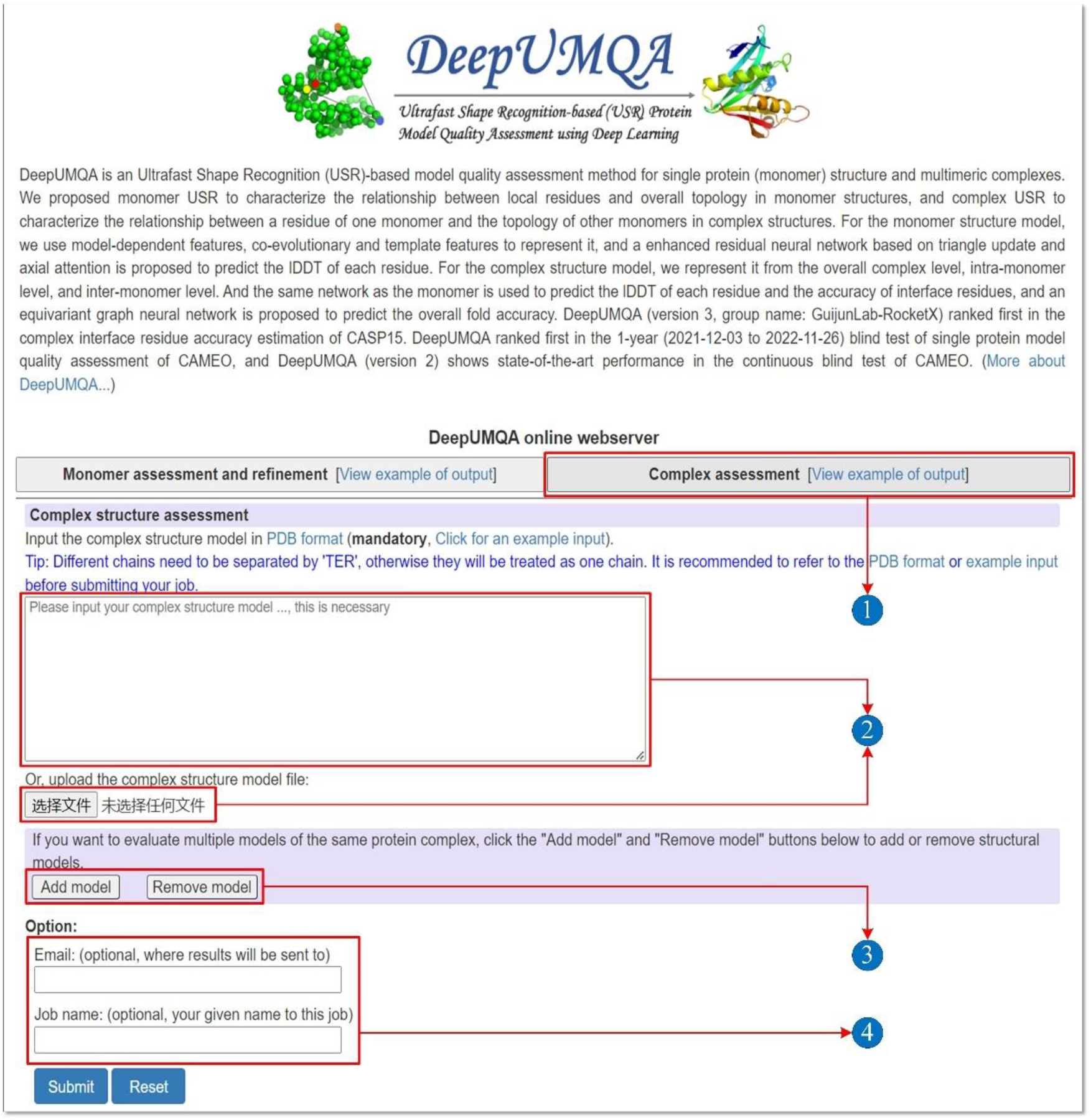
DeepUMQA3 server job submission. The DeepUMQA3 server is integrated into the DeepUMQA server, and the submission of complex model quality assessment tasks can be started by clicking the “Complex assessment” button (1). Users can enter the PDB of the complex structure through the text box or upload the PDB file (2). Via the “Add model” and “Remove model” buttons, multiple complex structures can be selected for evaluation (3). Users can optionally provide an “Email” to receive result notifications and a “Job name” (4). Submit or reset tasks via the “Submit” or “Reset” buttons.

**Figure S7.**
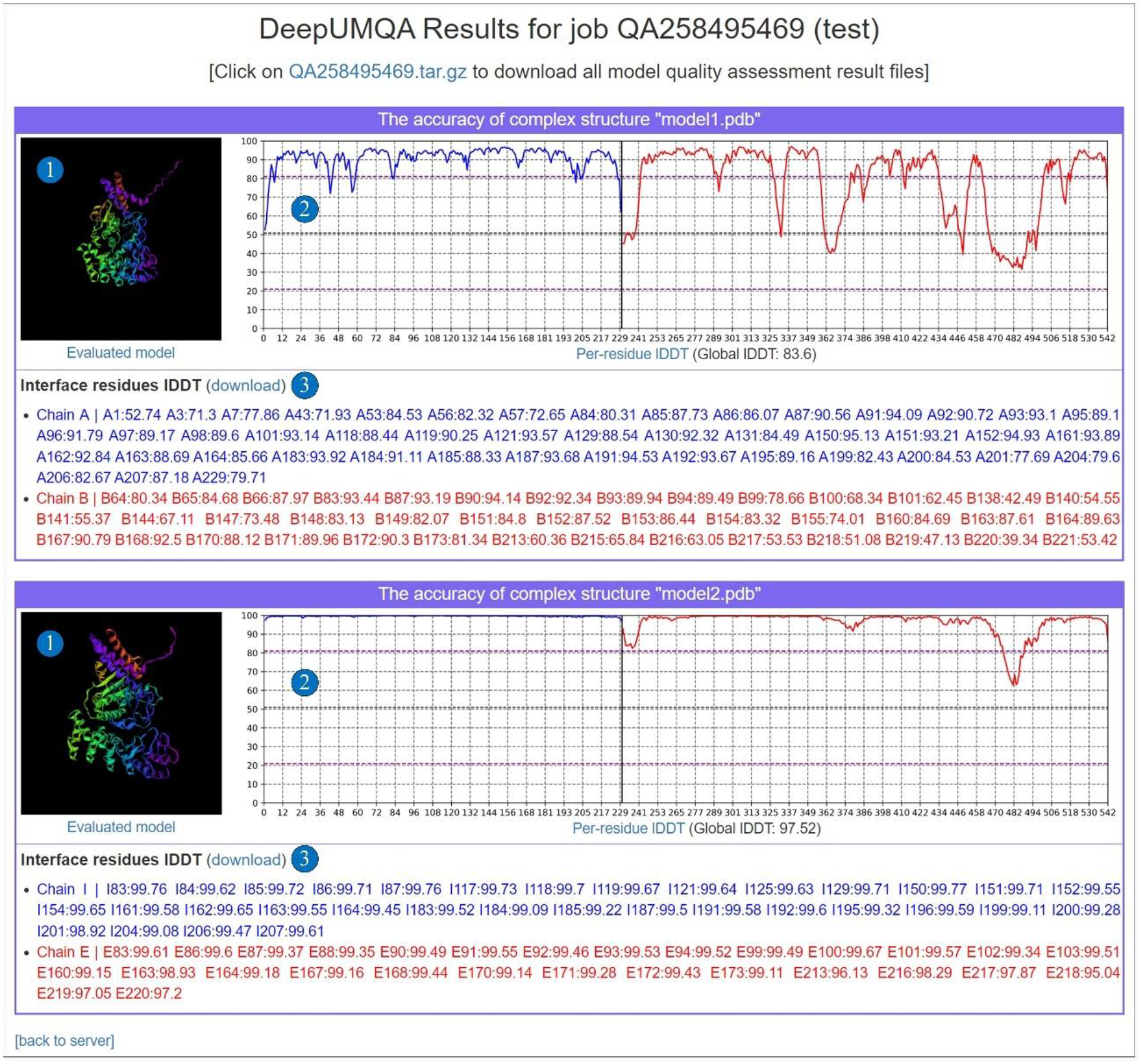
Example of DeepUMQA3 result web page. For each complex structure submitted by the user, the result web page will display the 3D structure (1), a graph of the per-residue lDDT (2), and the accuracy of the interface residues for each chain (3). Users can download each result individually, or download a compressed package of all results.

